# Phase of neural oscillations as a reference frame for attention-based routing in visual cortex

**DOI:** 10.1101/2021.11.08.467673

**Authors:** Ehsan Aboutorabi, Sonia Baloni Ray, Daniel Kaping, Farhad Shahbazi, Stefan Treue, Moein Esghaei

**Affiliations:** Department of Physics, Isfahan University of Technology, Isfahan, Iran; Indraprastha Institute of Information Technology, New Delhi, India; Helmholtz Centre for Environmental Research - UFZ, Leipzig, Germany; Cognitive Neuroscience Lab, German Primate Center, Goettingen, Germany

**Author notes:** Corresponding author: Moein Esghaei.

## Abstract

Selective attention allows the brain to efficiently process the image projected onto the retina, selectively focusing neural processing resources on behaviorally relevant visual information. While previous studies have documented the crucial role of the action potential rate of single neurons in relaying such information, little is known about how the activity of single neurons relative to their neighboring network contributes to the efficient representation of attended stimuli and transmission of this information to downstream areas. Here, we show in the dorsal visual pathway of monkeys (medial superior temporal (MST) area) that neurons fire spikes preferentially at a specific phase of the ongoing population beta (~20 Hz) oscillations of the surrounding local network. This preferred spiking phase shifts towards a later phase when monkeys selectively attend towards (rather than away from) the receptive field of the neuron. This shift of the locking phase is positively correlated with the speed at which animals report a visual change. Furthermore, our computational modelling suggests that neural networks can manipulate the preferred phase of coupling by imposing differential synaptic delays on postsynaptic potentials. This distinction between the locking phase of neurons activated by the spatially attended stimulus vs. that of neurons activated by the unattended stimulus, may enable the neural system to discriminate relevant from irrelevant sensory inputs and consequently filter out distracting stimuli information by aligning the spikes which convey relevant/irrelevant information to distinct phases linked to periods of better/worse perceptual sensitivity for higher cortices. This strategy may be used to reserve the narrow windows of highest perceptual efficacy to the processing of the most behaviorally relevant information, ensuring highly efficient responses to attended sensory events.

## Introduction

Spike counts and patterns are used to represent sensory information in neural systems [1–6]. Independent of the magnitude of spike rate, the temporal structure of the individual spikes relative to the collective activity of neighboring population of neurons conveys information about the sensory stimuli. It has been shown that the phase of the oscillatory neuronal population activities where the spikes occur at, carries information in the neural systems of a wide range of animal models [7,8]. For example, Montemurro et al. previously reported that in the primary visual cortex (i.e., V1 of anaesthetized monkey), the phase of neuronal spikes (relative to low-frequency oscillations) provides additional information, exceeding the information that can be extracted from the spike rate alone [8]. Recent research has provided evidence for the significance of the interaction between the spike times and the oscillatory phase in the ongoing population activity in association with high-level brain functions, such as visual attention [9–16], sensory processing [17,18], decision making [19], and memory [20–24]. In addition to information encoding [24–28], this spike-phase coupling, as an indication of spike timing control, mediates the inter-neuronal synchronization of activity within a region, that is of particular importance for not only optimizing the inter-areal communication via the temporal coordination of postsynaptic potentials [29–31], but also the performance of subjects’ perceptual decisions [32,33]. There have been increasing evidence suggesting that the brain uses the phase of oscillatory population activities to distinguish information items via separate phase alignment of spikes. By shifting the phase relationship between the spike trains and the underlying oscillatory population activity in a systematic way, known as the phase shift phenomenon, the brain classifies spikes according to their information. Data from rodents indicate that spatial information is encoded at specific phases of the ongoing population oscillations for the hippocampal pyramidal cells [27,34,35]. Similarly, prefrontal neurons encode different memory contents at distinct phases, allowing the brain to store and separate different objects in memory concurrently [24]. This strategy may be exploited by the visual cortex to differentiate behavioral relevance of different information items.

Due to the brain’s limited resources to process the environmental information, selective information processing is crucial for an efficient processing of behaviorally relevant information, a cognitive function of the mammalian brain through which some aspects of environment gain advantage over nearby distractors, known as selective attention [36–39]. Attention has been shown to affect visual neurons which represent the attended location/feature in a systematic way to enhance the neural representation and consequently the perception of behaviorally relevant (over irrelevant) sensory inputs which benefits behavior [40–54]. More recently, several studies have suggested a temporal structure for spatial/feature-based attention, i.e., attention samples the visual environment rhythmically, that is the brain rhythms govern the visual target’s detection performance [55–63]. This rhythmic sampling is an indication of alternating periods of better/worse perceptual sensitivity (assigned to ‘good’ and ‘poor’ phase of collective neural activity, respectively) [57,61,64].

Here we hypothesize that the phase of ongoing population oscillations may provide an internal reference frame for discriminating sensory information of different behavioral relevance (see [21] for consistent observations for working memory). We conjecture that the preferred phase of spiking may be used as reference when encoding stimuli at the attended location to allow for a selective routing to downstream areas.

To test this hypothesis, we recorded from the medial superior temporal cortex (MST), an area in the extrastriate visual cortex of monkeys, with neurons selective to spiral motion patterns [65,66]. We simultaneously recorded local field potentials (as a proxy of the collective synaptic activity of local neural populations [67–69]) and single cell activity from two rhesus monkeys engaged in a visual attention task. We compared the neurons’ preferred spiking phase when they were encoding relevant vs. irrelevant sensory information. Our data and the results of our computational modelling support our hypothesis that the temporal pattern of single neuronal spikes is shaped by the phase of beta oscillations in the LFP, and importantly, spatial attention organizes spike times in order to discriminate relevant from irrelevant sensory inputs by aligning the spikes to different phases of beta oscillations. These results provide insight into how the information of attended stimuli, rather than unattended stimuli might be preferentially read out by downstream brain areas.

## Results

It has been shown that single neurons spike at specific phases of their neighboring network’s activity. This spike-phase coupling is indicative of a neuron’s activity relative to its surrounding network. Here, we hypothesize that the location/feature towards which attention is allocated may be encoded by the coupling of the spikes’ timing to different phases of the local network’s oscillatory activity (Fig 1).

**Fig 1.**
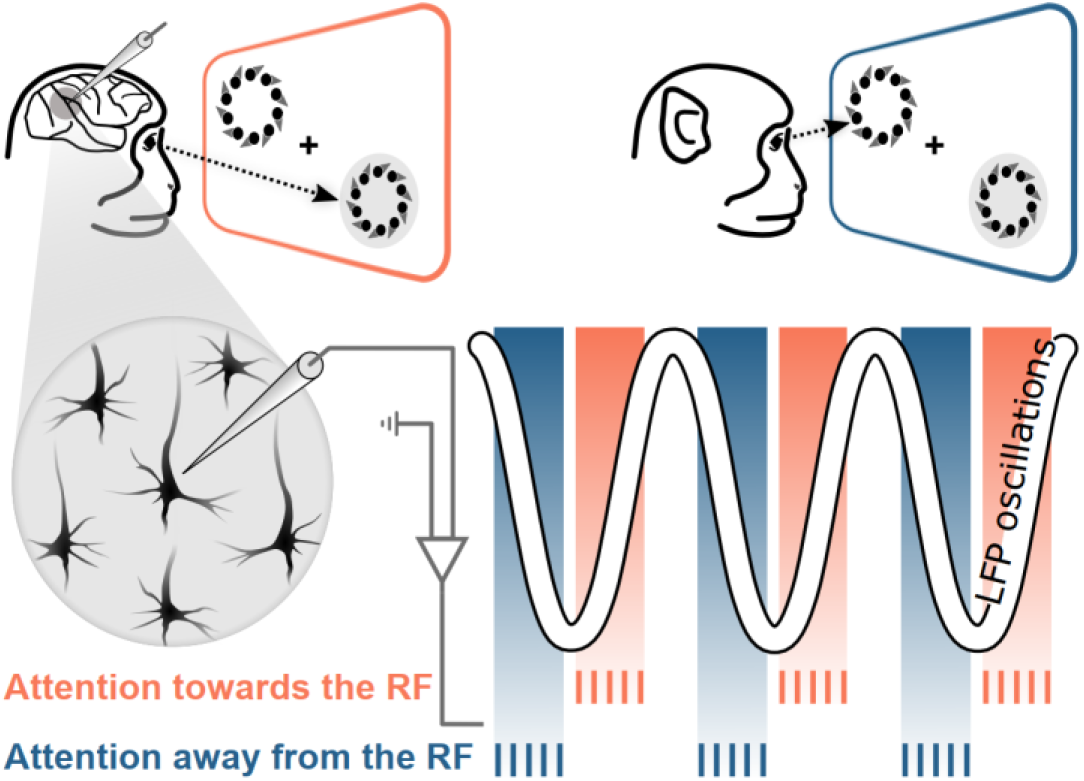
Phase coding of selective attention. We hypothesize that the activity corresponding to different positions/features in the visual field dominantly occurs at distinct instantaneous phases of the local oscillations in the visual cortex’s population activity. This implies that shifting the focus of attention from one position/feature to another position/feature shifts the timing of neuronal spikes between the two corresponding phase alignments. Therefore, changes in similarity between the focus of attention to the position/feature to which the sensory neuron is tuned, shifts the preferred spiking phase. Here, attention towards vs. away from the receptive field of a given neuron is depicted by different colors (orange and blue). Note, that the choice of preferred phase depicted here for the two conditions is arbitrarily selected.

To evaluate our hypothesis, we trained two rhesus monkeys to perform a visual attention task, where they had to direct their attention towards either of the two spiral motion patterns (SMP) presented on the screen. Monkeys N and W correctly performed the task successfully in 91% and 89% of those trials where they maintained their eye fixation. We recorded local field potentials (LFP) and single unit activity from 90 motion-selective neurons in visual cortical area MST of the monkeys while they performed the task. Depending on which of the two stimuli was cued and its direction of motion relative to the recorded neuron’s properties, trials came from one of three different attention conditions; They either attended to the preferred stimulus inside the receptive field (RF) (attend-in pref), preferred stimulus outside the RF (attend-out pref), or the anti-preferred stimulus outside the RF (attend-out null) (note that in all conditions, the stimulus inside the RF is the preferred stimulus) (Fig 2A). We compared the single unit responses between two of these attentional conditions (attend-in (Pref) versus attend-out (Null)) which differ in the attended stimulus’ location and direction, while the stimulus inside the RF is kept the same (preferred). Spike rate was significantly higher when attention was directed to the RF and the preferred stimulus, compared to when attention was directed outside the RF and the null stimulus. This confirmed that the animals did indeed selectively attend to the cued stimulus (Fig 2B). All of the following analyses focus on the spiking activity and LFP during the 500 ms interval starting from 350 ms following the onset of the SMPs. This allowed us to address the sustained component of the neural activity.

**Fig 2.**
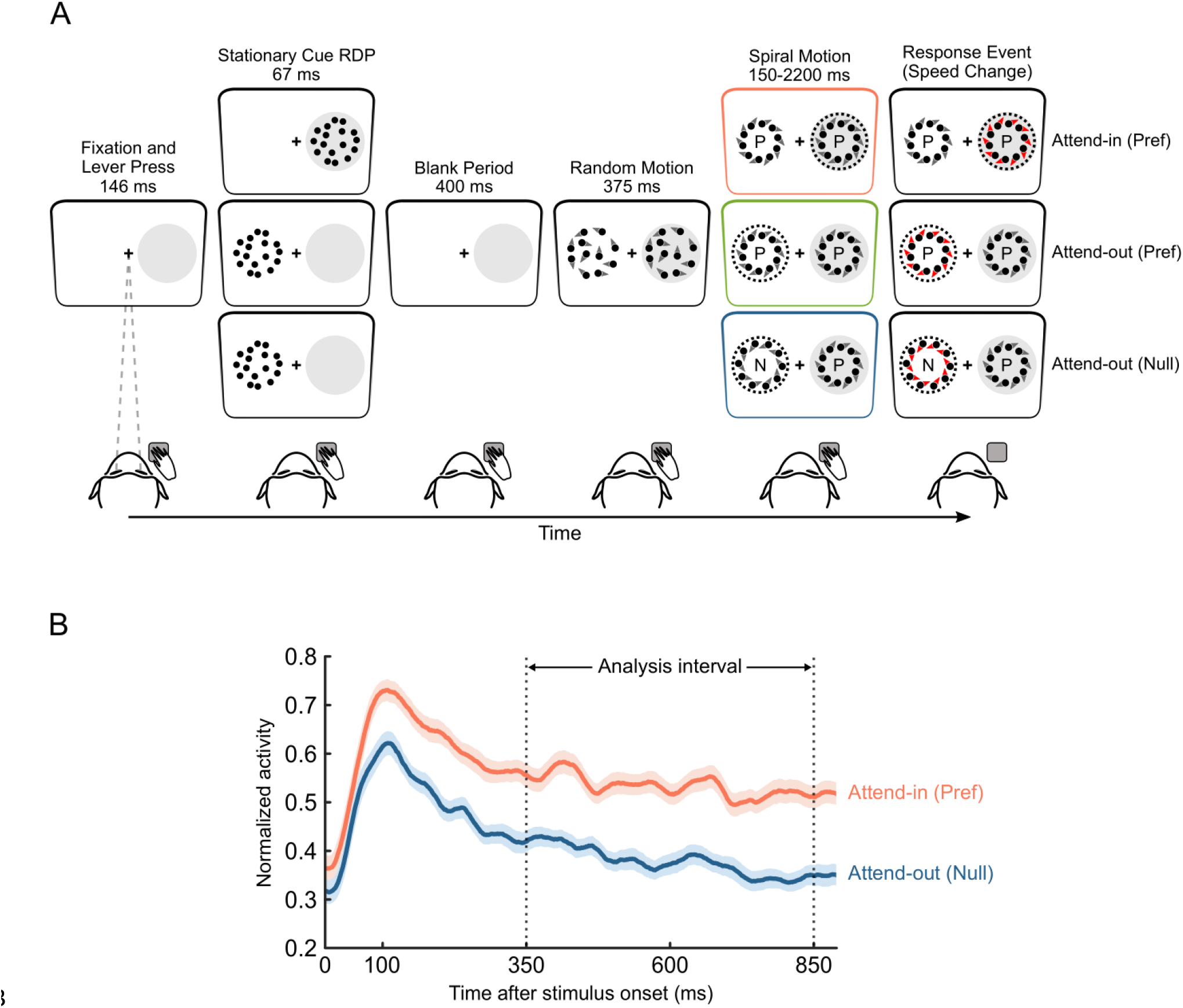
Behavioral task to study selective attention. (A) Schematic of the trial evolution for three attentional conditions. After the monkey fixed its gaze on the central fixation point and pressed a lever, an eccentric spatial cue (static random dot pattern) was presented to guide the animal’s spatial focus of attention. After a blank period, two randomly moving RDP masks were displayed at an equal eccentricity (one inside and the other outside the RF (gray-filled circle)), followed by two spiral motion patterns (SMP) moving in the preferred direction (here: expansion motion) or the null direction (contraction motion). Monkeys were rewarded when they responded to a brief speed increment in the target stimulus (indicated by a dashed circle here) happening at a random instance between 150-2200 ms. They further had to ignore the change in the distractor. The different types of attentional conditions are shown by the colors orange (attend-in(pref)), green (attend-out(pref)) and blue (attend-out(null)). (B) Normalized spike density functions across MST neurons (calculated with Gaussian kernel with σ = 15*ms*) for the attend-in (Pref) and attend-out (Null) conditions. Time 0 indicates the onset of SMPs and the vertical dotted lines indicate the analysis time range. Error bands represent the standard error of the mean (SEM).

To investigate if the focus of attention is associated with the phase of the local network where the spikes preferably occur at, we calculated the phase of LFPs (as a representative of the local population activity) where the spikes were preferably locked to (see S1 Fig for an illustration of phase-dependent firing rate for recorded neurons) and compared it between the two extreme attention conditions. Fig 3A shows the frequency-resolved difference of the preferred LFP phase between trials when attention was directed towards the inside of the neuron’s RF (preferred stimulus) or the outside of the RF (null stimulus). We found that switching attention towards the RF and the preferred stimulus was associated with a significant positive phase shift of beta oscillations (19-24 Hz, paired Watson-Williams test, p < 0.05, false discovery rate [FDR] corrected). This shift of attention focus did not have a significant effect on the strength of spike-phase coupling within the afore-mentioned frequencies (permutation test, p > 0.64 for all frequency bands, corrected for multiple comparisons-see Methods), suggesting that the beta frequency range is specifically involved in phase coding of neuronal spikes and not modulating the synchrony between neurons [10,70]. Fig 3A (right panel) plots the distribution of the preferred phases (0 denotes each neuron’s preferred phase averaged across conditions) across neurons for the two attentional conditions at a sample frequency point (highlighted in yellow, corresponding to 18-22 Hz). These results indicate that the two different attentional conditions are associated to different phases of the beta range. This phase shift between the two extreme attentional conditions suggests that different locations and/or features of the attended stimulus may be encoded by different phases of LFPs. The phase shift predominantly reflects the effect of spatial attention (paired Watson-Williams test, p < 0.05, FDR corrected), rather than feature-based attention (paired Watson-Williams test, p > 0.52, FDR corrected), as illustrated in S2 Fig. We further focused on neurons with a higher effect of attention (see S3 Fig for neurons with a minor effect of attention on firing rate). An attentional index was computed by dividing the difference in average activity between the two attention conditions by their sum (see materials and methods). Similar analysis as Fig 3A was performed for neurons with a large enough attentional index (>0.1) (Fig 3B-left panel), showing that the attentional phase shift is larger for neurons demonstrating a stronger effect of attention on their spike rate (0.80 and 0.08 rad for AI>0.1 and AI<0.1, respectively; Watson-Williams test over the average of frequency bands shown as significant in panel A, p = 0.017). Fig 3B (right panel) presents the neuron-wise phase shift distribution averaged across frequency bands with a significant phase shift (highlighted in cyan).

**Fig 3.**
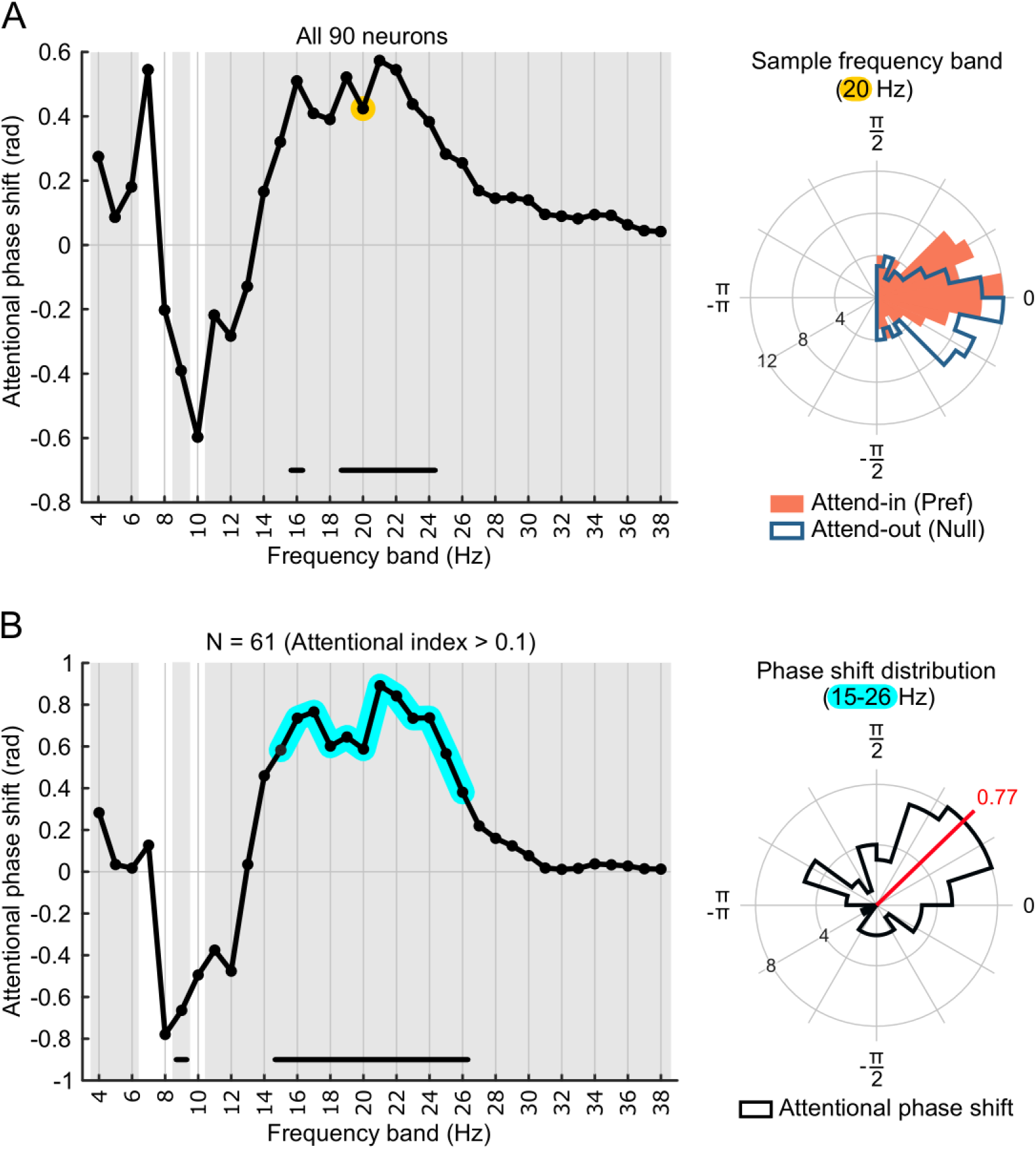
Focus of attention determines the preferred LFP phase of spikes. (A) The left panel shows the difference in the preferred phase of spikes between the two attention conditions (attend in-Pref – attend out-Null) across different frequency bands (for 4 Hz-wide bands sliding by 1 Hz) (see S4 Fig for a similar analysis for each of the two animals). There is a significant shift of the preferred LFP phase for the frequency bands indicated by the horizontal bar (Watson-Williams test, p < 0.05, FDR corrected). The X-axis indicates the middle frequency of each band and the Y-axis represents the preferred phase’s shift between the attention conditions. The frequency bands with a non-uniform distribution in their phase shift are indicated using a gray background (Rayleigh test for circular uniformity on the phase shift distribution, p < 0.05, FDR corrected for multiple comparisons). The right panel plots the distribution of preferred phase for the attend-in (Pref) trials and attend-out (Null) trials for a sample frequency band (highlighted in yellow on panel left). The average preferred phase of each neuron is aligned to 0 degree here. (B) The left panel illustrates the attentional phase shift across different frequency bands for neurons with an attentional index of at least 0.1. Right panel presents the distribution of the average attentional phase shift across the 12 frequency bands with a positive and significant phase shift (highlighted in cyan). The red line indicates the circular average.

We next asked if the magnitude of the phase shift is linked to the behavioral response of monkeys. To this end, we examined the relationship between the attentional phase shift and the monkeys’ reaction time (RT) in detecting the target change. The trials were first sorted into four quartiles based on their RT and the phase shift was calculated based on the first and fourth quartile (short RT and long RT, respectively). Fig 4 shows that the attentional phase shift is only significant for trials with the shorter RT within the beta range. This confirms that the attentional phase shift is tightly linked to the speed of responses. The association of phase to behavior is observed in both attend-in(Pref) and attend-out(Null) trials (S5 Fig).

**Fig 4.**
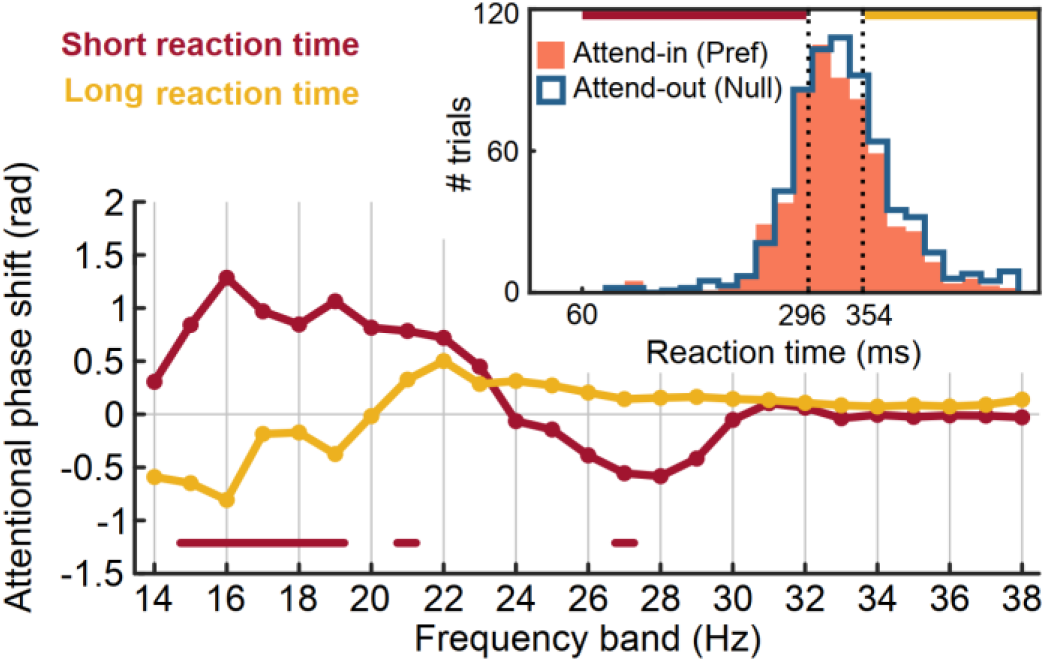
Reaction time (RT) varies as a function of phase shift. The observed phase shift within the beta range is mainly observed for trials with a short RT (fast behavioral responses). Inset: distribution of RTs in each attentional condition. The first (red) and fourth (yellow) quartiles are referred to as short RT and long RT, respectively.

To investigate the underlying mechanisms of attentional phase coding, we established a computational model. The model is constructed of two internally connected neural populations, representative of the sensory and regulatory subnetworks (Fig 5), each of which consisting of excitatory and inhibitory neurons. Each of the model’s neurons could receive excitatory/inhibitory postsynaptic potentials from neurons of the same or the other subnetwork. Furthermore, external drives to the model neurons were adjusted so that the two subnetworks showed oscillatory activity predominantly at frequencies around 40 and 20 Hz (for the sensory and regulatory network, respectively - see Methods). We next examined whether applying different delay levels to the postsynaptic potentials could influence the coupling of sensory neurons to the modulatory subnetwork’s beta rhythm. We observed that, by increasing the synaptic delay, sensory neurons start to shift their preferred spiking phase relative to modulatory beta oscillations (Fig 5A), presumably due to a delayed effect of the modulatory subnetwork on the sensory neurons. These results suggest that the neural mechanism whereby attention shifts the preferred phase of neurons, may involve controlling the synaptic delay of inter-neuronal connections within a sensory area.

**Fig 5.**
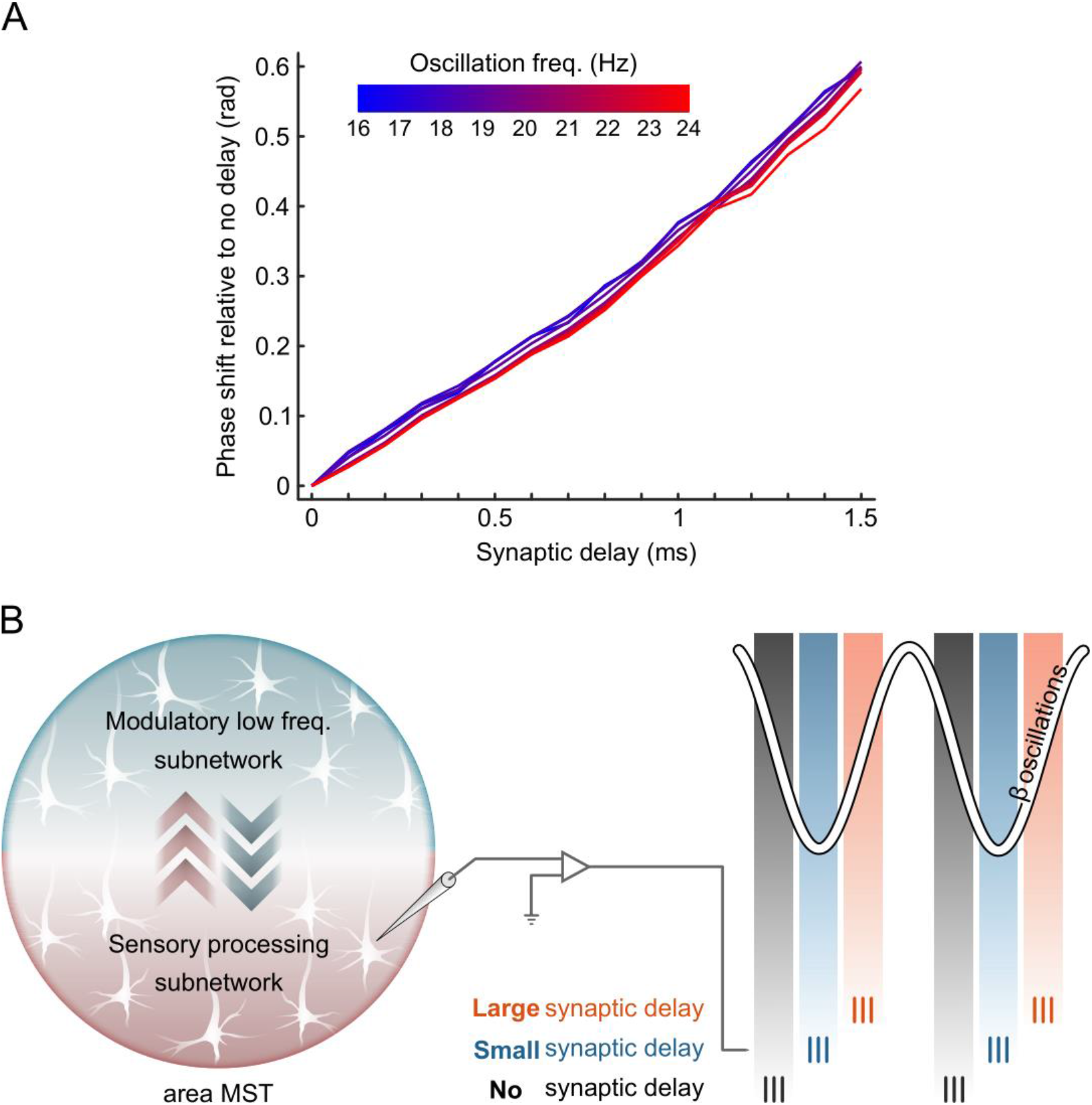
Synaptic delay of postsynaptic potentials adjusts the timing of spikes relative to the phase of the modulatory network. (A) Our computational model reveals that the predominant phase of spiking activity of sensory neurons relative to the population’s beta oscillations is correlated with synaptic delay. The X-axis represents the synaptic delay applied to the timing of postsynaptic potentials and the Y-axis represents the phase of coupling relative to that when there is no synaptic delay. (B) This schematic illustrates the model’s output showing that neurons may couple to different phases at different synaptic delays. We speculate that attention may manipulate the preferred phase of sensory neurons’ spiking activity relative to an intra-areal subnetwork via imposing synaptic delay to postsynaptic potentials.

Our data suggest that the focus of attention is associated to the timing of the activity of MST neurons relative to the phase of the neighboring neural network’s pattern of activation. We further found that the magnitude of this phase difference enforced by shifts of attention focus is linked to the behavioral output of the subjects. These are supportive of the account that the preferred phase of neuronal firing relative to the neighboring network is an indicator of the location towards which the subjects attend.

## Discussion

Here, we investigated whether the temporal structure of the discharges of single neurons relative to the phase of oscillatory activities among their neighboring neuronal network (captured by the LFP) plays a functional role in relaying the attended stimulus’ information. Our data reveal that first, neuronal spikes are locked to a specific phase of beta oscillations of the LFP in the macaque visual cortical area MST. Second, spatial attention modulates this preferred phase of the locking and that this modulation is especially pronounced for neurons strongly modulated by attention. Third, the attentional shift of the preferred beta phase is predictive of the subsequent behavioral response time of animals. Our computational model shows that this attentional shift of the preferred phase may be associated to a so-called attentional lag of post-synaptic potentials within the network of sensory areas.

In our study we observed that spikes preferentially cluster at different phases of the ongoing beta rhythm of LFP oscillations depending on the spatial focus of attention. This observation suggests that attended/unattended sensory information are channeled by different phases of beta oscillations (Fig 6). Specifically, we observed that with the shift of spatial attention from the outside to the inside of the RFs, spikes tend to fire at a later phase of the beta oscillations (Fig 3A). Given the independence of a neuron’s firing *rate* from its firing *phase*, this feature of the MST network allows downstream areas to decode the locus of attention from sensory neurons even if they do not exhibit an attention-dependent firing rate modulation. The neural mechanism for reading out of the focus of attention in downstream areas is a question for future studies.

**Fig 6.**
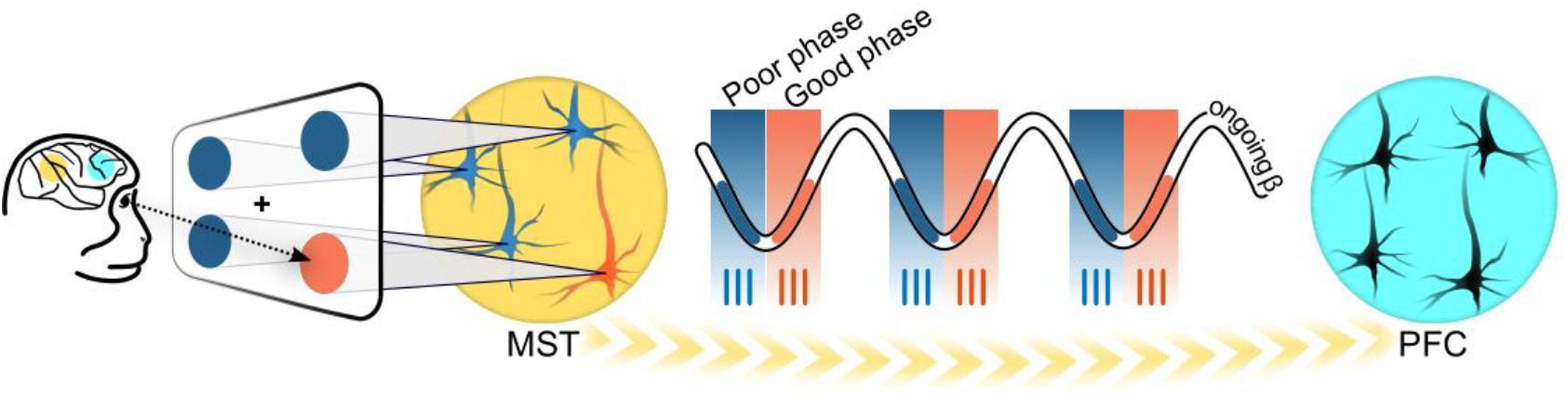
Network oscillations as an internal reference frame for distinguishing spikes encoding attended vs. unattended stimuli. This schematic illustrates the speculative mechanism that allows attended sensory information to be transmitted from area MST to a higher cortical area (such as PFC) at an optimal phase of beta optimizing relevant information’s readout. MST neurons whose receptive fields coincide with the attended location (orange neuron) fire action potentials at an optimal phase (‘good phase’) of ongoing beta oscillations of LFP which can facilitate the readout mechanism in the higher-order area. But other neurons with receptive fields outside the locus of attention (blue neurons) are activated at a different phase linked to worse neuronal sensitivity in higher area (‘poor’ phase). In this way attended visual information could be processed approximately in a way as if unattended information is absent.

In light of previous studies which have shown that, unlike unattended information which may not be perceived consciously at all, the attended sensory inputs are processed more efficiently and rapidly [40,41], our results are indicative of the existence of functionally different beta phases; a more informative beta-dependent phase (‘good’ phase, where spikes are locked to when attention is shifted towards the RF of the neuron, indicative of better information coding, (Fiebelkorn et al., 2018)), where the MST neurons encode the attended information and transmit it efficiently to higher order cortices. Whereas the spikes encoding the less-attended irrelevant stimulus, occur at the ‘poor’ beta phase, less likely to reach downstream cortical areas. (Fig 6).

Our data further indicate that the attentional phase shift is linked to behavioral performance, i.e., the faster the animals respond the larger the attentional phase shift we observe. A large attentional phase shift can be interpreted as a sharper distinction of the attended and unattended information, helping to encode the attended information more efficiently by assigning the limited processing epochs to the most relevant stimuli in the scene. A sufficiently large attentional phase shift allows the attended stimulus to be selectively transmitted to higher cortical areas and read out with minimal distortion by the surrounding noise (the unattended stimuli).

Unlike the classic view suggesting that spatial attention samples the visual space continuously, several recent studies have revealed a rhythmic sampling by spatial attention [56,57,59,61,64,71]. Consistently, our results also indicate that MST neurons transmit the information of the attended stimulus to higher cortices through oscillatory beta-aligned periods of enhanced coding for the attended stimulus (Fig 6). Future studies need to examine the existence of a beta-aligned perceptual sensitivity for motion recognition.

One could interpret location in visual space as a feature in the visual feature space. Similarly, a neuron’s receptive field could be interpreted as the neuron’s preferred feature in location-space. Therefore, both types of attention modulate neurons’ responses based on the similarity of the attended feature relative to the neuron’s preferred feature [38,48]. Therefore, one might wonder whether the neural activity influence exerted by spatial and feature-based attention may have identical encoding mechanisms. However, this is in contrast to our finding regarding the modulation of firing phase, which demonstrates the neuronal modulation of locking phase is exclusively caused by spatial attention. Spatial attention takes advantage of firing phase through an exclusive mechanism to enable the visual system to selectively process the attended stimulus in more efficient manner. This supports the hypothesis that spatial and feature-based attention do not use the same mechanisms. Spatial attention modulates both firing rate and the preferred firing phase of neurons, whereas feature-based attention is not observed to modify the preferred firing phase.

One question is whether attentional phase coding follows a topographic map or not, i.e., if the distance between an attended and unattended stimulus (proportional to the distance between the RFs of the neurons representing them) is correlated with the magnitude of the attentional phase shift. This is particularly important when asking if beta phase can serve as an internal reference frame for distinguishing the attended stimulus from other stimuli. This could not be addressed in the current study due to lack of enough data and remains a question for future studies.

Together, our data show that action potentials in the macaque visual cortical area MST are locked to the phase of beta oscillations in the LFP and spatial attention significantly shifts this locking phase. We suggest that attention changes the locking phase to transmit relevant visual information through exclusive periods linked to higher perceptual sensitivity for higher cortices.

## Materials and methods

Research with non-human primates represents a small but indispensable component of neuroscience research. The scientists in this study are aware and are committed to the great responsibility they have in ensuring the best possible science with the least possible harm to the animals [72].

Methodological details concerning our animal subjects, their holding and welfare, our experimental permits, surgeries and implants as well as details of the experimental setup were reported previously [16,70,73–75]. We reiterate relevant details here:

### Subjects and animal welfare

Data were collected from two male rhesus monkeys (Macaca mulatta, Monkey W, Monkey N, both 12-year-old males). Area MST was accessible through a recording chamber implanted over the parietal lobe based on a magnetic resonance imaging (MRI) scan (right hemisphere for Monkey W, left hemisphere for Monkey N). Each monkey was implanted with a titanium head holder to minimize head movements during the experiment. Both monkeys were seated in custom-made primate chairs and head-fixated during the experiment.

All animal procedures of this study were approved by the responsible regional government office (Niedersaechsisches Landesamt fuer Verbraucherschutz und Lebensmittelsicherheit (LAVES)) under the permit number 33.14.42502-04-064/07 and were performed in full accordance with relevant guidelines and regulations.

The animals were pair-housed in the facilities of the German Primate Center (DPZ) in Goettingen, Germany. The facility provides the animals with an enriched environment (incl. a multitude of toys and wooden structures [76,77], natural as well as artificial light, exceeding the size requirements of the European regulations, including access to outdoor space. The animals’ psychological and veterinary welfare was monitored by the DPZ’s staff veterinarians, the animal facility staff and the lab’s scientists, all specialized in working with non-human primates. During the study the animals had unrestricted access to food and fluid, except on the days where data were collected or the animal was trained on the behavioral paradigm. On these days, the animals were allowed access to fluid through their performance in the behavioral paradigm. Here the animals received fluid rewards for every correctly performed trial.

Surgeries were performed aseptically under gas anesthesia using standard techniques, including appropriate peri-surgical analgesia and monitoring to minimize potential suffering. The two animals were healthy at the conclusion of our study and were subsequently used in other studies.

We have established a comprehensive set of measures to ensure that the severity of our experimental procedures falls into the category of mild to moderate, according to the severity categorization of Annex VIII of the European Union’s directive 2010/63/EU on the protection of animals used for scientific purposes (see also [78]).

### Behavioral task and electrophysiological recordings

The two monkeys were trained to perform a visual attention task (Fig 2A) in which they foveated a central fixation point (0.2°×0.2°; degrees of visual angle) and initiated each trial by pressing and holding a response lever. After 146 ms, a spatial cue (a stationary random dot pattern (RDP-4°large, dot density of 20 dots/degree)) appeared for 67 ms at the location where the animal had to attend covertly. Afterwards, the cue disappeared, and following a 400ms blank period, two mask RDPs were presented on the right and the left side of the screen. One of these mask RDPs was placed in the recorded neuron’s RF while the other one was placed outside the RF at an equal eccentricity from the fixation point. After 375 ms, the mask RDP at the neuron’s RF was replaced by a Spiral Motion Pattern stimulus (SMP) evoking a maximal neuronal response (preferred stimulus) and the mask RDP located outside the RF at the opposite hemi-field was replaced by a preferred/null SMP (null stimulus: an SMP evoking the minimal neuronal response). Importantly, the stimulus inside the RF always moved towards the preferred pattern of the neuron. The stimuli at the cued and uncued location are referred to as ‘target’ (designated by a dashed circle in Fig 2A) and ‘distractor’, respectively. At a random instant between 150-2200 ms after the onset of SMPs, a brief speed increment occurred either in the target or distractor stimulus and the animal was rewarded by receiving a drop of fluid after reporting the speed change in the target (and not in the distractor) by releasing the response lever. Depending on the location and motion pattern of the target stimulus, there were three attentional conditions; a. Attend-in (Pref), where attention was directed towards the receptive field (RF) of the recorded neuron, with a stimulus moving towards the preferred direction of the neuron. b. Attend-out (Pref), where attention was directed away from the RF of the recorded neuron, with a stimulus moving towards the preferred direction of the neuron. c. Attend-out (Null), where attention was directed outside the receptive field (RF) of the recorded neuron, where a null stimulus was presented. Therefore we were able to study the effect of spatial and feature-based attention by comparing the first two and the last two conditions, respectively.

Experiments were performed in a dim room and the procedure for stimulus presentation, monitoring the eye position and recording neural and behavioral data during the experiment were controlled by a custom computer program running on an Apple Macintosh PowerPC. We monitored the eye position using a video-based eye tracking (ET49, Thomas Recording, Giessen, Germany). Visual stimuli appeared on the computer monitor (at a distance of 54 cm and covered 40°×30° of visual angle at a resolution of 40 pixel/deg).

Here, we recorded the activity of 40 MST neurons from monkey N and 50 single-units from monkey W while they were performing the above-mentioned visual attention task.

### Data analysis

For all analyses, we used all correctly performed (hit) trials from the two animals and the data were analyzed using customized scripts in MATLAB, as well as the EEGLAB and CircStat toolboxes.

For all statistical tests, alpha level for the decision criterion was chosen to be 0.05 and to overcome the problems due to multiple testings, we used the false discovery rate (FDR) approach to determine adjusted p-values (based on the number of performed tests).

### Spike Rate quantification

To estimate changes in spike rate over time, the spike density function (SDF) was computed by convolving the spike train from each trial with a Gaussian kernel of SD=15 ms. For each unit, SDFs were averaged across trials of the same attentional condition and normalized to the maximum value across the two conditions’ average SDFs. The SDFs began to diverge between the two attentional conditions 67 ms after the stimulus onset (FDR adjusted p-value < 0.05). By focusing on the time window between 350-850 ms after the stimulus onset and including the trials where the speed change occurred following this interval, we ensured to focus only on the sustained rather than the transient part of the neural responses.

### Attentional Index measurement

To assess attention’s effect on spike rate, we calculated the attentional index (AI) for each recorded neuron within the interval of 350-850 ms after stimulus onset by comparing neuronal responses in the two different attentional conditions using the following formula:

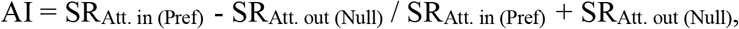

where SR_Att. in (Pref)_ and SR_Att. out (Null)_ are the mean spike rate when attention was directed towards the preferred stimulus in the RF or towards the null stimulus outside the RF, respectively.

### Spike-LFP phase coupling

For each recorded spike-LFP pair, we first band-pass filtered the LFP signals in 35 overlapping frequency bands with the bandwidth of 4 Hz and starting frequencies from 2 to 36 Hz (using eegfilt function from EEGLAB toolbox with an order of 3*sampling_rate/low_cutoff_freq) and then measured the similarity of phases where the spikes coincided with, at different frequency bands by defining the complex number z that is equivalent to the ‘power-biased circular sample mean’ of the phase distribution of spikes which occurred during the associated trial. In a single trial, the z vector (at different frequency bands of LFP) was calculated using the following equation:

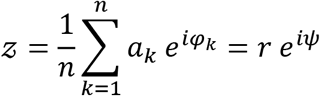

where *φ*_*k*_ and *a*_*k*_ represent the instantaneous phase and power of the filtered LFP, respectively determined by the argument and absolute value of Hilbert transform at the instant that the *k*th spike occurred and n is the total number of spikes occurring during each trial. The factor *a*_*k*_ weights the spike phases proportional to the instantaneous power, allowing spike phases measured with a higher signal to noise to proportionally have a large effect on the resultant vector. The complex number z can also be rewritten by parameters r and *ψ* (known as Kuramoto order parameters) which capture the strength of coupling and the locking phase, respectively. At a constant power, r is higher when spikes tend towards an identical phase and lower when spikes are uniformly distributed across the phase space. At each frequency band we obtained two z vectors (corresponding to the two attention conditions) for each neuron by calculating the circular average of the z vectors across trials (weighted by the average power of the filtered LFP within each trial) of each attention condition. To do so, we utilized the following equation:

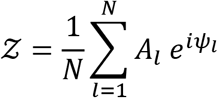

where *ψ*_*l*_ and *A*_*l*_ are the z vector’s phase and the filtered LFP’s average power (respectively) corresponding to the *l*th trial, and N is the neuron’s number of trials in each attend-in/attend-out condition.

By subtracting the phases of z vectors between attention conditions (attend-in (Pref) – attend-out (Null)) we determined the attentional phase shift and then by taking the absolute values of z vectors we estimated the PLVs assigned to the two attention conditions for each neuron. Therefore, we had 35 phase shift distributions for 35 frequency bands and by calculating the circular mean of each of them we determined a mean attentional phase shift within each frequency band (Fig 3). We examined for the similarity of the phase shift across neurons, by applying a Rayleigh test for non-uniformity of the phase shift distributions in each frequency band (gray filled areas in Fig 3 indicate frequency bands in which the phase shift data is sampled from a von Mises distribution). Afterwards we examined each of the frequency bands to see if there was a phase shift distribution with a mean unequal to zero by applying a paired Watson-Williams test (as a one-way ANOVA test for circular data).

### Modeling

Each of the subnetworks used to construct intra-regional neuronal circuits (i.e., sensory and regulatory subnetworks) is composed of 200 Hodgkin-Huxley-like model neurons, 80% excitatory (*N*_*E*_) and 20% inhibitory (*N*_*I*_). The probability of establishing excitatory and inhibitory projections (*P*_*E*_ and *P*_*I*_, within/between subnetworks) are 0.3 and 0.2, respectively.

Our excitatory and inhibitory model neurons are of the form of the Hodgkin-Huxley model [79], but with different parameter values and time rate functions. The membrane potential *V* and ionic gating variables evolve according to the following:

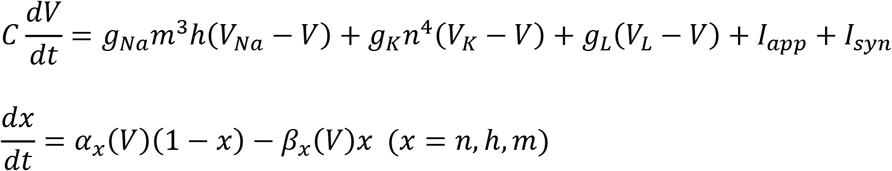

where *C*, *V*, *t*, *g* and *I* denote capacitance density (*μF*/*cm*^2^), voltage (*mV*), time (*ms*), ionic conductance density (*mS*/*cm*^2^) and current density (*μA*/*cm*^2^), respectively. *α* and *β* are functions of the membrane potential (measured in *ms*^−1^) and for gating variable *m* we used the following assumption: *m* = *m*_∞_(*V*) = *α*_*m*_(*V*)/[*α*_*m*_(*V*) + *β*_*m*_(*V*)].

The constants and time rate functions of our excitatory neurons are [80]:

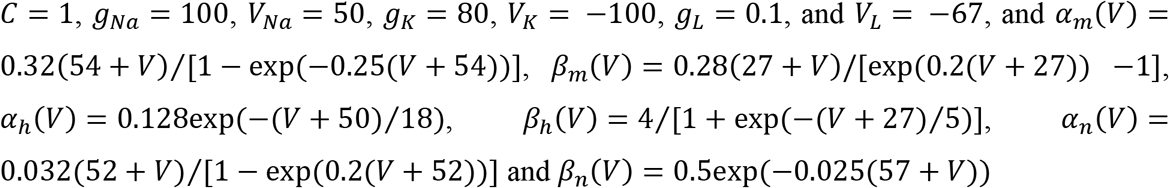

whereas the constants and time rate functions of our inhibitory neurons are [80,81]:

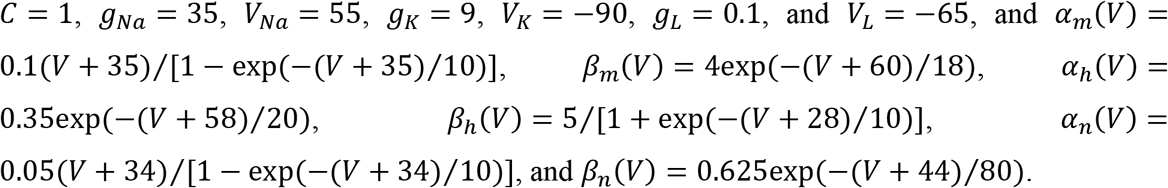

The first subnetwork (i.e., the regulatory subnetwork) is driven at an around 20 Hz rhythm, while the other (i.e., the sensory subnetwork) is driven at around 40 Hz, by applying a stationary heterogeneous external excitatory drive *I*_*app*_ to the excitatory cells, chosen from a Gaussian distribution with a variance of 0.05 and mean values 0.12 and 0.4, respectively.

Excitatory synaptic input to each neuron is modeled by a term of the form *I*_*syn*_ = *g*_*E*_(*V*_*rev,E*_ − *V*) Σ *s*_*i*_, where *g*_*E*_ = 0.0019/*P*_*E*_*N*_*E*_ is a Gaussian variable with the coefficient of variation 20%, and *V*_*rev,E*_ = 0. The sum extends over those excitatory cells that send projections and the synaptic gating variable *s* is defined in the form [82]:

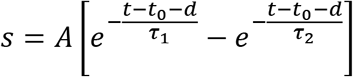

where *t*_0_, *d* and *A* are the presynaptic spike time, synaptic delay and normalization factor respectively, and τ_1_ = 3 ms and τ_2_ = 1 ms determine the rise and decay synaptic time constants. Inhibitory synaptic input to each neuron is modeled by a term of the form *I*_*syn*_ = *g*_*I*_(*V*_*rev,I*_ − *V*) Σ *s*_*i*_, where *g*_*I*_ = 0.01/*P*_*I*_*N*_*I*_ is a Gaussian variable with a coefficient of variation 5%, and *V*_*rev,I*_ = −75. The sum extends over those inhibitory cells that send projections and the synaptic gating variable *s* is the same as that for excitatory synaptic inputs, however here we have τ_1_ = 4 ms and τ_2_ = 1 ms.

The system of differential equations was solved in MATLAB using the midpoint method with time step Δ*t* = 0.01 ms. The system was initialized asynchronously [83], i.e., we chose the initial state of neurons driven above their spiking threshold in a way that in the absence of synaptic currents, it will take each neuron a time span of *uT* to fire its first spike, where *T* is the intrinsic spiking period and *u* ∈ [0,1] is a uniformly-distributed random variable, and the remaining neurons (those driven below their spiking threshold) are initialized close to their stable state (on their phase plane). The system was solved for 500,000 steps (i.e., 5000 ms) and as a convention, spike times were taken to be the moments at which the membrane potential crosses −20 mV from above. Here, we considered the population-averaged membrane potential as a heuristic of the collective neuronal activity, a computational analogue of the LFP which is called LFP-like signal [68,82] and successfully captures the information content of the recorded LFP, although it fails to reproduce the recorded spectrum [68].

To examine if there exists any systematic relationship between synaptic delay and the preferred phase of the spiking activity, we considered the synaptic delay *d* as a Gaussian-distributed random variable with a variance of 0.05 ms. We then increased its mean in the range of 0 to 1.5 ms by steps of 0.1 ms, carried out the simulation and finally calculated the preferred spiking phase of the pooled spike times of the sensory subnetwork relative to beta oscillations of the regulatory neuronal population (Fig 5A).

## Acknowledgments

The project was supported by the Deutsche Forschungsgemeinschaft (DFG, German Research Foundation) – Projektnummer 154113120 – SFB 889, project C04. We thank Leonore Burchardt, Sina Plümer, Klaus Heisig, Dirk Prüsse, and Ralf Brockhausen for skilled technical assistance and Beatrix Glaser for her administrative support.

## Data availability statement

The data underlying the result figures are publicly available on figshare (https://doi.org/10.6084/m9.figshare.16940899).

## Supporting information

**S1 Fig.**
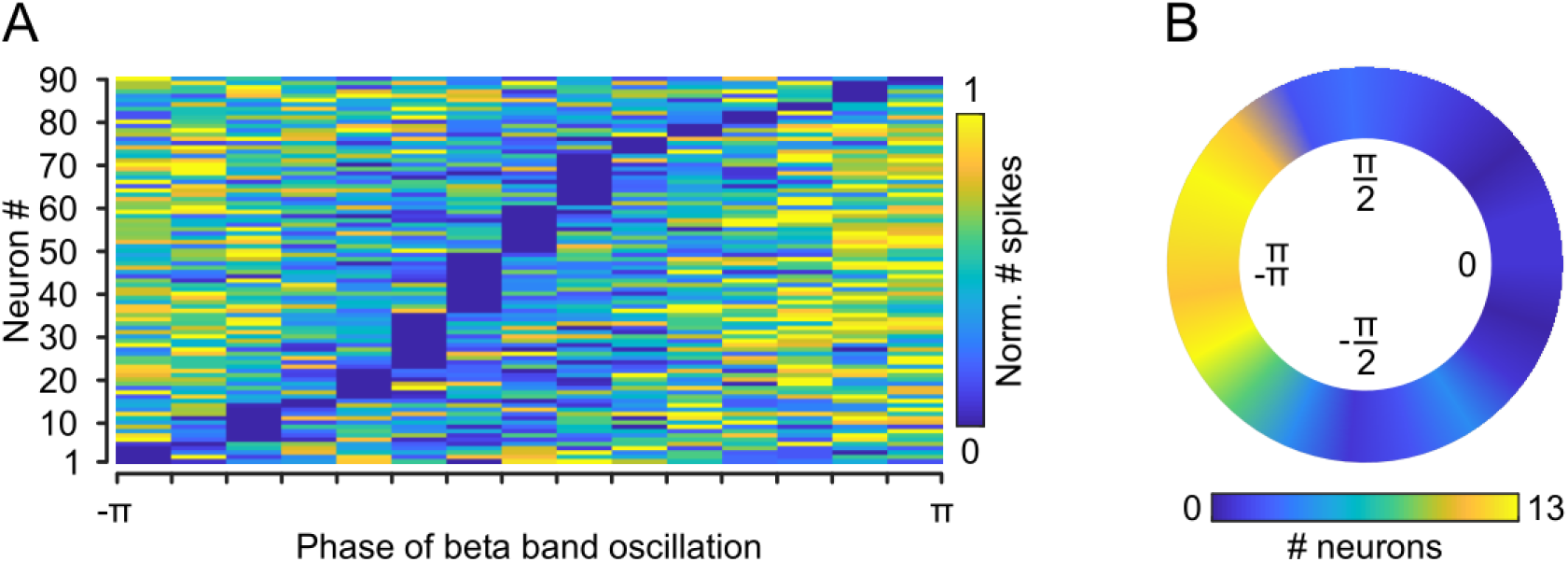
MST neurons exhibit a phase-dependent firing pattern. (A) We computed the density of spikes within different phase bins (bin-size = 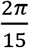) of the beta band (sample frequency range of 21-25 Hz) oscillatory activity and normalized it between 0-1 for each trial. For each condition we averaged the phase-dependent density distributions across the corresponding trials and next, computed the mean distribution across the two conditions for each neuron. This Fig shows the normalized (within the 0-1 range) distribution for each neuron. Neurons are sorted based on the phase with the smallest spike density (Y-axis). (B) Color-plot presents the distribution of the preferred spiking phase and indicates that the MST neurons tend to fire around the trough of beta oscillations (Rayleigh test for circular uniformity, p<10^−9^). For each neuron we calculated the preferred spiking phase relative to beta oscillations by averaging the neurons’ locking phase across the two attentional conditions (see METHODS) and then averaging the resultant phase across beta bands within which the attentional phase shift was significant (see Fig 3A). The phase resolution is based on a division of the phase spectrum into 19 non-overlapping bins.

**S2 Fig.**
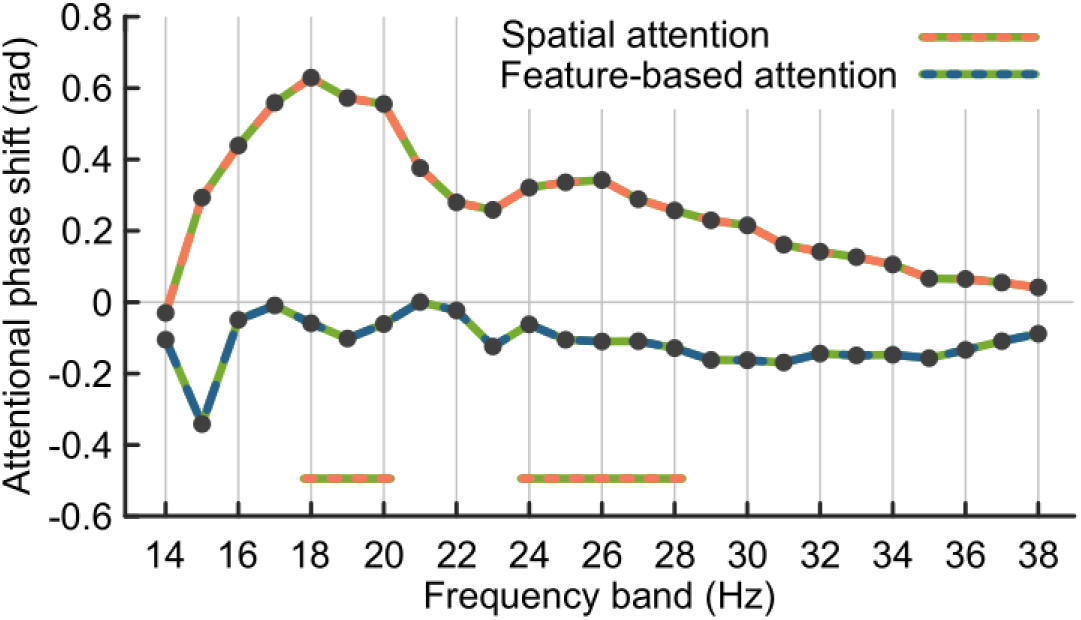
Attentional phase shift is dominated by spatial, rather than feature-based attention. The blue-green curve (for feature-based attention) shows the phase difference between attend-out (pref) and attend-out (null), whereas the orange-green curve (for spatial attention) presents the phase difference between attend-in (pref) and attend-out (pref) conditions. The bold horizontal lines indicate the frequencies with a significant attentional phase shift.

**S3 Fig.**
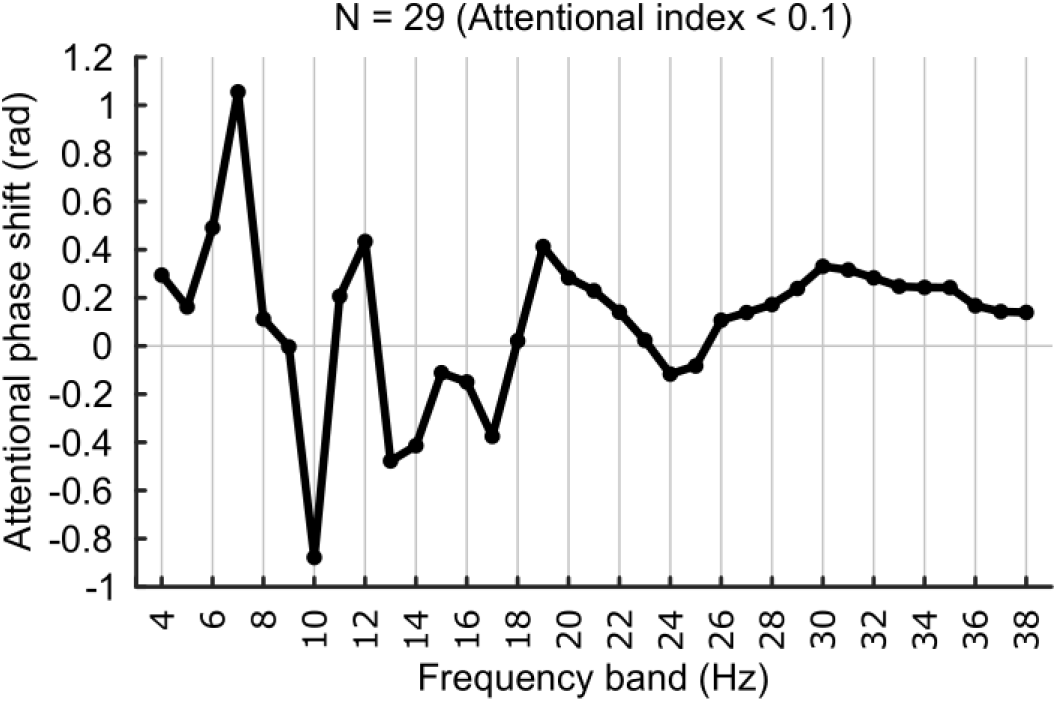
Absence of a significant attentional phase shift for neurons with an AI < 0.1 (minor effect of attention).

**S4 Fig.**
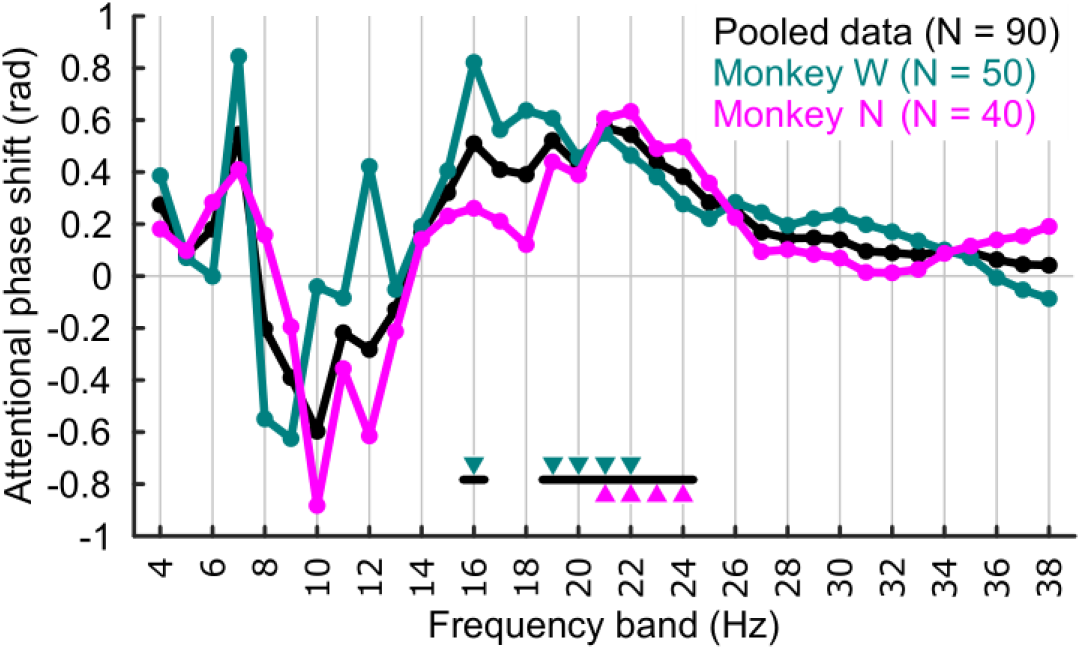
Frequency-resolved attentional phase shift for each animal. We performed similar analysis as of Fig 3A, separately for each monkey. The frequency bands showing a significant phase shift among the frequencies which showed a significant phase shift for the pooled data of both animals (black horizontal bars), are indicated for each monkey’s data using the triangle with the corresponding color.

**S5 Fig.**
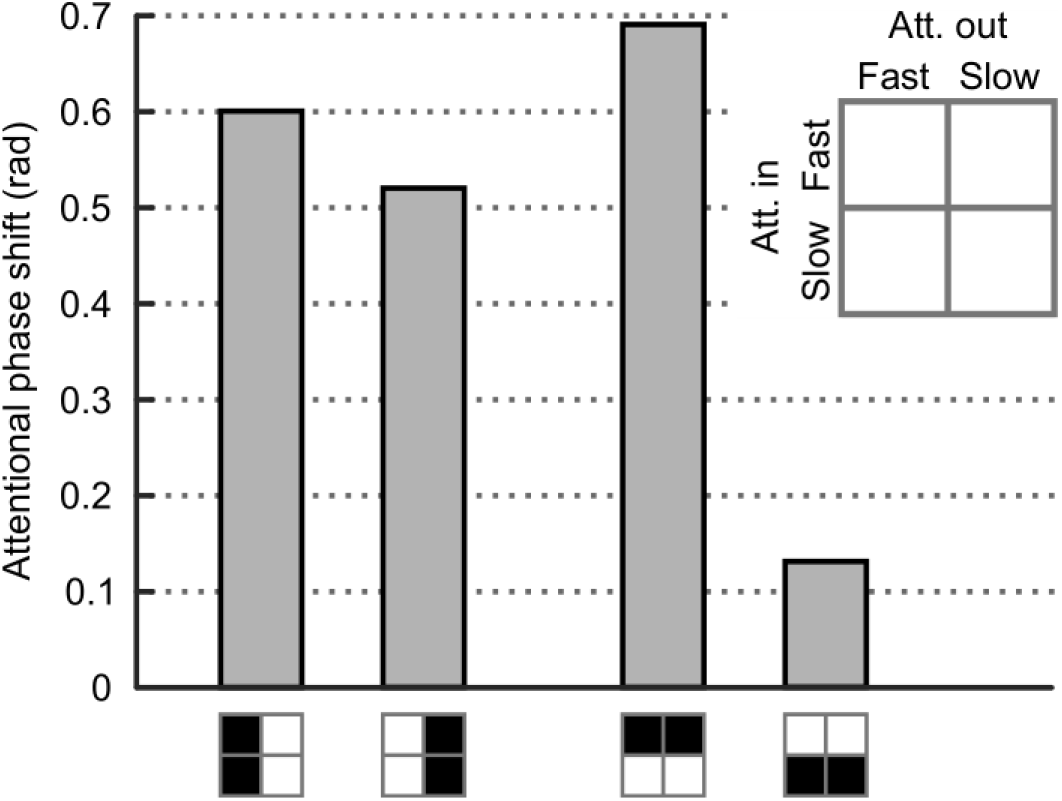
Magnitude of the phase shift is linked to the behavioral response in both attention conditions. In each condition we ascendingly sorted the trials based on the monkey’s RT and then considered the first and fourth quartiles as fast and slow trials, respectively. By using four possible combinations of these sets of trials (shown by different colors), we calculated the phase shifts for each neuron in each frequency band using the corresponding trials in each pair of sets. Averaging across frequency bands, a phase shift distribution (across neurons) was yielded for each combination of trial sets. The height of each bar in the above Fig indicates the circular-mean of the phase shift distribution which was formed by merging two distributions. X-axis illustrates the specific pair of trial sets pooled in this analysis. The relationship between fast and slow trials is independent of if they are extracted for the attend-in (the right-most pair of bars) or attend-out (the left-most pair of bars) trials. These results indicate that the association of phase to behavior is observed in both attend-in and attend-out trials.

